# Reconstructing unobserved cellular states from paired single-cell lineage tracing and transcriptomics data

**DOI:** 10.1101/2021.05.28.446021

**Authors:** Khalil Ouardini, Romain Lopez, Matthew G. Jones, Sebastian Prillo, Richard Zhang, Michael I. Jordan, Nir Yosef

**Affiliations:** Department of Electrical Engineering and Computer Sciences, University of California, Berkeley, USA; École CentraleSupélec, Gif-sur-Yvette, France; École Normale Supérieure Paris-Saclay, Gif-sur-Yvette, France; Department of Cellular and Molecular Pharmacology, University of California, San Francisco, USA; Biological and Medical Informatics Graduate Program, University of California, Berkeley, USA; Department of Statistics, University of California, Berkeley, USA; Center for Computational Biology, University of California, Berkeley, Berkeley, USA; Ragon Institute of MGH, MIT and Harvard, USA; Chan Zuckerberg Biohub, San Francisco, USA

## Abstract

Novel experimental assays now simultaneously measure lineage relationships and transcriptomic states from single cells, thanks to CRISPR/Cas9-based genome engineering. These multimodal measurements allow researchers not only to build comprehensive phylogenetic models relating all cells but also infer transcriptomic determinants of consequential subclonal behavior. The gene expression data, however, is limited to cells that are currently present (“leaves” of the phylogeny). As a consequence, researchers cannot form hypotheses about unobserved, or “ancestral”, states that gave rise to the observed population. To address this, we introduce TreeVAE: a probabilistic framework for estimating ancestral transcriptional states. TreeVAE uses a variational autoencoder (VAE) to model the observed transcriptomic data while accounting for the phylogenetic relationships between cells. Using simulations, we demonstrate that TreeVAE outperforms benchmarks in reconstructing ancestral states on several metrics. TreeVAE also provides a measure of uncertainty, which we demonstrate to correlate well with its prediction accuracy. This estimate therefore potentially provides a data-driven way to estimate how far back in the ancestor chain predictions could be made. Finally, using real data from lung cancer metastasis, we show that accounting for phylogenetic relationship between cells improves goodness of fit. Together, TreeVAE provides a principled framework for reconstructing unobserved cellular states from single cell lineage tracing data.

## Introduction

Recent advances in CRISPR/Cas9-based genome engineering and single-cell sequencing assays have enabled the simultaneous measurement of lineage information and transcriptomic state at the single-cell resolution [1, 2]. Already, these technologies have been been used to study mammalian embryogenesis [2] and cancer metastasis [3], amongst other applications. In these studies, researchers use phylogenies - tree structures that describe relationships between observed cells - to infer transcriptomic determinants of dynamic phylogenetic patterns. A key limitation thus far, however, derives from the fact that only the leaves of these trees are directly observed whereas the internal nodes of the tree represent unobserved, *ancestral* states. While rich insights can be yielded from the relationships between leaves alone, accurately inferring these ancestral states of a tree would allow researchers to formulate far more sophisticated “evolutionary” models of biological processes [4].

The variational autoencoder (VAE) [5] is a powerful framework for fitting flexible generative models to data in a scalable fashion. In particular, such tools have been applied in several areas of molecular biology [6], including modeling of single-cell RNA sequencing (scRNA-seq) data. Most of the subsequent research focuses on the case where each datapoint is an independent replicate of the same generative process, providing better variational distributions or more flexible models [7, 8, 9]. In this work, we seek to exploit the tree structure generated from lineage tracing as prior information about the sample-sample covariance structure. In doing so, we seek to fit a prescribed model that can be used to predict ancestral expression across the tree in a principled fashion. Here we introduce TreeVAE, a fully-probabilistic approach that builds on previous work, such as the time-marginalized coalescent VAE [10], by tailoring the inference to bespoke observation models for scRNA-seq and rich phylogenies inferred from CRISPR/Cas9-based lineage tracing data. After describing our generative model and an inference procedure for it (Section 1), we compare TreeVAE to alternative methods on simulated and real datasets (Section 2).

## 1 Tree Variational Auto-encoder (TreeVAE)

### 1.1 Formal Description of the Generative Model

We assume that we know a phylogeny 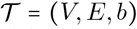, a directed rooted tree with vertex set *V*, edge set *E* and edge length function *b*. We note the weight *b*(*e*) of an edge *e* = (*u,v*) as *b*_u,v_. The vertex set *V* is partitioned into *L* leaf vertices *V_L_* = {1,…, *L*} (a set of cells) and *I* internal vertices *V_I_* = {*L* + 1,…, *L* + *I*} (ancestral cell states) such that *V* = *V_L_* ∪ *V_I_*. Phylogenies may be inferred from single-cell lineage tracing mutation data using publicly available methods such as Neighbor Joining [11] or Cassiopeia [12].

We introduce a probabilistic model describing the evolution of latent random variables *z_v_* at every vertex *v* along the phylogeny 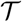. Those latent variables correspond to a low-dimensional embedding of cell states, as explored in previous work [13]. Unlike in most applications of generative models to single-cell data, where those variables are independently sampled for every cell, we model correlation between cells with a Gaussian Random Walk (GRW) on 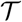. For the root node *r*, we sample from an isotropic Gaussian distribution:

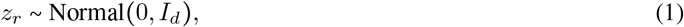

where *d* denotes the dimension of *z_r_*. Then, every vertex *v* ∊ *V* \ {*r*}, *z_v_* is sampled according to an isotropic Gaussian distribution centered at its parent’s location and with covariance scaled by the edge length:

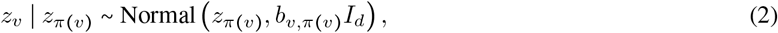

where *π*(*v*) is the unique parent of *v* in 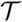.

We observe the sc**RNA**-seq measurements *x*_1:*L*_ = {*x*_1_;… *x_L_*}, as well as the library size (i.e., the number of gene counts per cell) *α*_1:*L*_ = {*α*_1_,… *α_L_*} at each of the leaves. We propose to build on an observation model previously used for single-cell transcriptomics data [13] whereby for each cell *l* and for every gene *g*, the gene expression *x_lg_* is generated as:

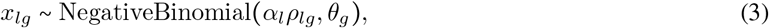

where *ρ_lg_* = *f^g^*(*z_l_*) is the output of a neural network (*f* has a softmax output layer for normalization purposes) and *θ_g_* is a gene-wise global parameter learned by variational Bayes.

This model assumes that the correlation between cells is fully-characterized by the GRW as illustrated in Figure 1 and described in the following likelihood decomposition:

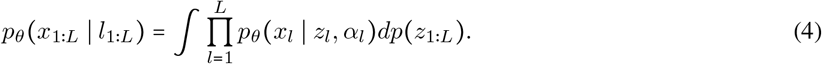

**Figure 1:**
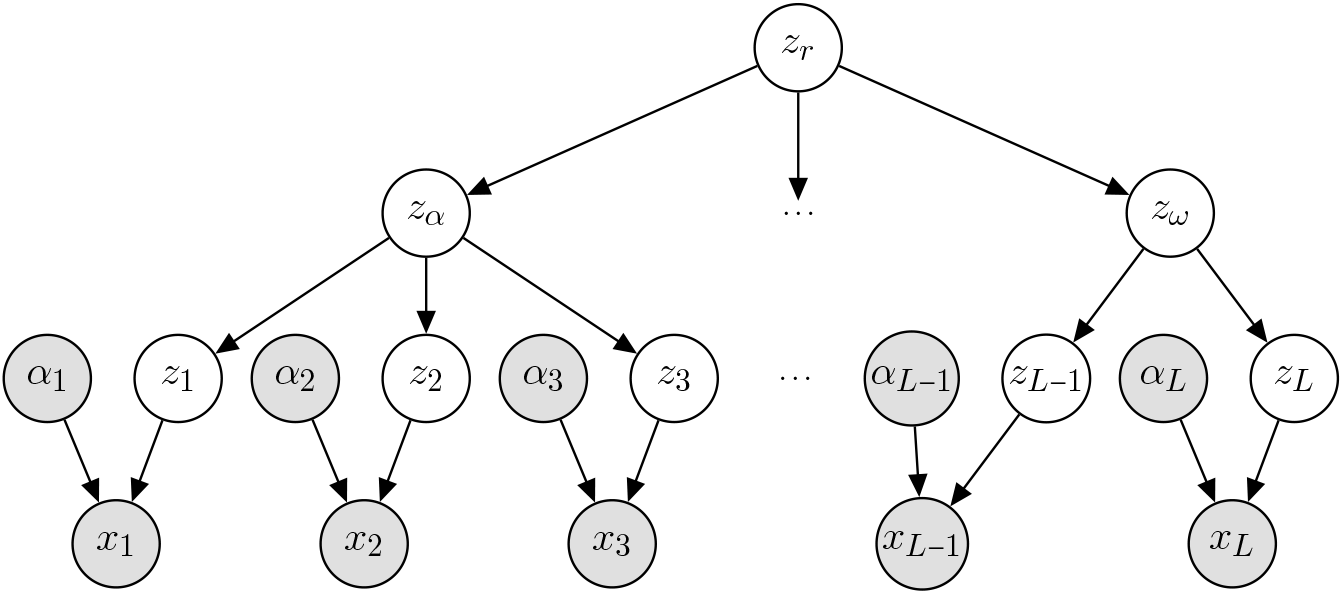
The proposed graphical model. Shaded vertices represent observed random variables. Empty vertices represent latent random variables. Edges signify conditional dependency.

This model is an extension of the SVAE [14], and is a particular instance of the TMC-VAE [10], for which the phylogeny is known *a priori*. We refer to those models as TreeVAEs in this manuscript.

### 1.2 Inference

As with standard VAEs, the marginal likelihood of (4) is intractable. We therefore develop a variational inference recipe to (i) learn the parameters *θ* of the model and (ii) approximate the posterior distribution *p*(*z*_1:*L*_ | *x*_1:*L*_). Because our model couples a complex non-linear observation model with a more simple correlation model in latent space (any marginal distribution for the GRW is tractable), we may build upon previous work (SVAE and TMC-VAE) to derive a variational inference recipe.

We introduce a mean-field variational approximation to the posterior *p*(*z*_1:*L*_ | *x*_1:*L*_) which we assume factorizes as: 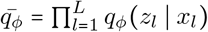, and we derive the evidence lower bound (ELBO):

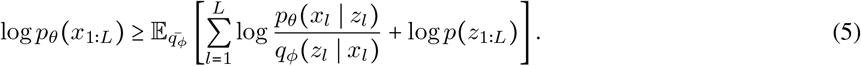

Provided that one can calculate the marginal likelihood *p*(*z*_1:*L*_) of the latent variables over the leaf nodes, and the gradient of the ELBO with respect to the parameters of the variational distribution, one may then learn the parameters of the generative model (*θ*) and the inference model (*ϕ*) via stochastic gradient ascent on the ELBO [5]. However, the complexity of naive calculations of this marginal distribution is cubic in the number of leaves, and therefore unsuitable. Fortunately, the marginalization of these latent variables is tractable in linear time, using a message passing algorithm on the tree [15]. Although usually those algorithms are only derived for bifurcating trees, we propose here an extension for multifurcating trees. Below, we derive the marginals for the base case of bifurcating trees (a triplet). We describe the general recursive algorithm in Appendix A.

#### Triplet example

Let (*x_r_, x_a_, x_b_*) be a triplet of random variables, following a local Gaussian diffusion model:

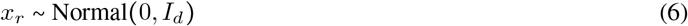

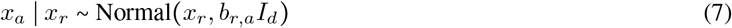

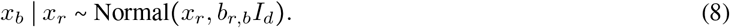

We note as *ϕ*(· *μ,σ*^2^) the probability density function of the multivariate Gaussian distribution with mean *μ* and covariance *σ*^2^*I*. Completing the square, we have that *p*(*x_a_, x_b_* | *x_r_*) identifies as an unnormalized Gaussian density:

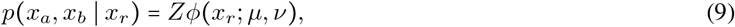

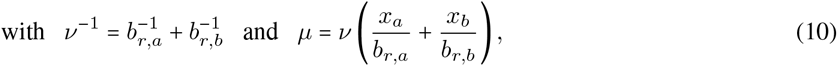

and a tractable normalization constant *Z*. The marginal likelihood *p*(*x_a_, x_b_*) simply follows from integrating out the prior *p*(*x_r_*).

### 1.3 Posterior Predictive Density

Unlike the vanilla VAE, our TreeVAE model explicitly helps define posterior predictive densities for latent variables *z_i_* and gene expression measurements *x_i_* at each internal node *i* ∊ *V_I_* of the phylogeny. On the latent space, the posterior predictive is approximated as:

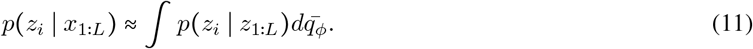

Similarly, on the feature space, the posterior predictive is approximated as:

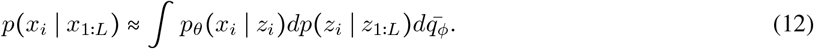

All quantities appearing in (11) and (12) are readily available after training, except for the conditional distribution *p*(*z_i_* | *z*_1:*L*_) for which we use another message passing algorithm, that also has linear time complexity (Appendix A).

## 2 Performance Benchmarks

To assess the performance of our approach, we evaluate the accuracy of the posterior densities. This is a sensible approach, as we seek to correctly estimate the cellular states at ancestral nodes of the phylogeny. Towards this end, we report several metrics to evaluate the quality of *p_θ_*(*z_i_ | x*_1:*L*_) (on latent space), as well as *p_θ_*(*x_i_* | *x*_1:*L*_) (on feature space). Because the ground truth for these densities is generally intractable, we propose three sets of experiments. First, we benchmark the quality of our approximate posterior predictive density on a Gaussian process factor analysis model (Section 2.1), for which the ground truth is tractable. Second, we provide a benchmark on a simulated single-cell RNA sequencing data, using the prior predictive density as (approximate) ground truth (Section 2.2). Finally, we explore the application of TreeVAE to real single-cell data from cancer metastasis (Section 2.3).

For the experiments in Section 2.1 and 2.2, we simulate ground truth tree topologies using a generalized birth-death model. A tree is simulated by beginning with a single node. We use two exponential distributions, parameterized by *α* and *β*, to model the time until a cell divides (i.e., birth of a new lineage) and the time until a cell dies, respectively. We repeat the birth-death process until a desired number of leaves is reached.

Throughout, we compare the imputation accuracy of TreeVAE to two baselines. The first one is a naive approach based on averaging gene expression at all the leaves beneath an internal node. The second one is based on averaging the latent space for leaves below an internal node from a fitted VAE before decoding. Such heuristics have been used in several scRNA-seq data analysis scenarios [16]. We ran VAE and TreeVAE on a NVIDIA TITAN Xp, and fitting the data took less than a few minutes for every dataset.

### 2.1 Gaussian Process Factor Analysis Simulations

As a first test-bed, we consider a simulation framework based on the Gaussian process factor analysis model [17], for which posterior predictive densities are tractable (fully-specified in Appendix B, with derivations for the ground truth). This model is obtained by using a linear Gaussian conditional distribution in place of the negative binomial decoder in (3):

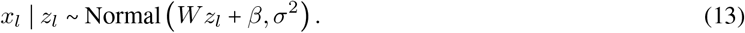

We adapt the observation model of the VAE and the TreeVAE accordingly. Each simulated dataset is based on a tree with 100 leaves (cells) and 100 genes. All results are averaged across ten simulations.

In this setting, we compare the posterior predictive on the latent variables *p_θ_(z_i_* | *x*_1:*L*_) as well as the one on the features *p_θ_*(*x_i_* | *x*_1:*L*_) to their respective ground truth. On the latent space, we report the mean square error of the mean of the posterior predictive *p_θ_*(*z_i_* | *x*_1:*L*_) (MSE; lower is better), the *k*-nearest neighbors purity (Purity; higher is better) [18] and the cross entropy between the prior distribution *p_θ_*(*z*_1:*L*_) and the approximate posterior 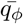 (CE; lower is better). (To note, latent space metrics do not apply for the average baseline.) On the feature space, we report the average Pearson correlation for imputation of ancestral gene expression across all genes (*r*; higher is better), as well as the Spearman correlation (*ρ*; higher is better), and the mean-square error for gene expression (MSE; lower is better).

We report the results in Table 1. On latent space, TreeVAE outperforms the VAE with respect to all three metrics. In particular, the MSE metric suggests that TreeVAE learned a more accurate posterior approximation compared to VAE. Moreover, the *k*-NN purity and cross entropy metric show that the latent space inferred by TreeVAE is more reflective of the phylogeny compared to VAE (as expected). On the feature space, TreeVAE also outperforms baselines on all three metrics. Although all the correlation scores are rather high for all methods in this setting, TreeVAE still provides a substantial improvement over the naive baselines.

**Table 1:** Results on the Gaussian process factor analysis simulations (averaged across ten different simulations).

|  | LATENT SPACE |  |  | FEATURE SPACE |  |  |
| --- | --- | --- | --- | --- | --- | --- |
| | MSE | Purity | CE | $r$ | $\rho$ | MSE |
| Average | - | - | - | 0.828 | 0.803 | 0.768 |
| VAE | 2.28 | 0.372 | 2,515 | 0.846 | 0.821 | 0.863 |
| TreeVAE | **1.89** | **0.450** | **281** | **0.869** | **0.844** | **0.541** |

### 2.2 Gaussian Process Poisson Log-normal Simulations

Although the linear Gaussian system described previously helps diagnose the effectiveness of our inference procedure, it remains an unrealistic model for describing scRNA-seq data. Therefore, we instead use a Poisson Log-normal observation model as a more realistic simulation framework. More precisely, in place of the observation model described in (3), we generate *x_l_* as:

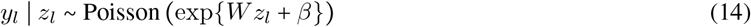

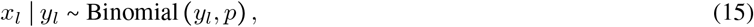

in which *p* was adjusted to bring the frequency of zeros to 80% in the dataset. Each simulated dataset is based on a tree with 500 leaves (cells) and 1000 genes. All results are averaged across ten simulations. Because the ground truth distribution for both posterior predictive densities are not accessible, we slightly modify the evaluation metrics presented previously. First, we do not report MSE on the latent space because it cannot be evaluated. Second, because the gene expression posterior predictive *p_θ_*(*x_i_* | *x*_1:*L*_) is no longer tractable, here we use the prior predictive distribution *p_θ_*(*x_i_*) as a proxy for ground truth. We report the results in Table 2. On the latent space, TreeVAE outperforms the VAE on both metrics. This emphasizes again that the TreeVAE produces representation at internal nodes that are more reflective of the tree structure. On the feature space, we observe that the correlation scores are generally low compared to the previous experiment, likely due to the high level of noise from binomial sub-sampling. However, even in this setting we observe similar trends as from our previous experiment: the TreeVAE outperforms both metrics. Finally, we investigate the relationship between certainty and accuracy in our TreeVAE model. As expected, our estimates become more uncertain closer to the root and this in turn affects model accuracy (Figure 2).

**Table 2:** Results on the Gaussian process Poisson Log-normal simulations (averaged across ten different simulations). MSE is reported on normalized counts.

|  | LATENT SPACE |  | FEATURE SPACE |  |  |
| --- | --- | --- | --- | --- | --- |
| | Purity | CE | $r$ | $\rho$ | MSE |
| Average | - | - | 0.350 | 0.314 | 7.53 |
| VAE | 0.523 | 8,481 | 0.402 | 0.324 | 5.81 |
| TreeVAE | **0.615** | **1,577** | **0.413** | **0.327** | **5.80** |

**Figure 2:**
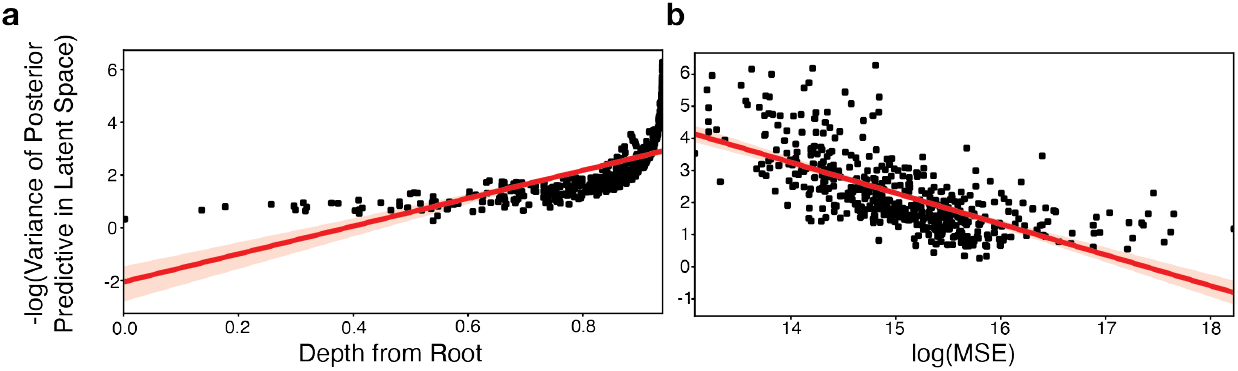
Relationship between uncertainty and error in the Gaussian Process Poisson Log-normal simulation experiments with the TreeVAE model. (a) Variance of posterior predictive density on latent space for each internal node compared to the depth (Pearson’s *r* = 0.6761). (b) Error and uncertainty of each prediction (Pearson’s *r* = −0.6765).

### 2.3 Analysis of Cancer Metastasis Data

We next assess the performance of the TreeVAE model on real CRISPR/Cas9 single-cell lineage tracing data. As a proof of concept, we analyzed a single clone of 603 cells from a recent dataset that traced the lineages of lung cancer tumors as they metastasized throughout a mouse [3]. Here, we leverage the tree reconstructed from CRISPR/Cas9 barcodes with Cassiopeia [12] as used in the original study. Because Cassiopeia does not explicitly model the edge weights of the phylogeny, we separately inferred these based on the assumption of an ultrametric tree. Finally, because a large fraction of genes detected by scRNA-seq do not have a strong relationship to the lineage, we only considered the top 100 genes autocorrelated with the phylogeny, as evaluated by *Hotspot* [19].

We fit the TreeVAE with the observation model described in (3). In this experiment, we have limited ground truth because neither latent variables nor gene expression is known at the ancestral nodes of the phylogeny. We still propose to compare to scVI [13], a VAE with identical observation model as the TreeVAE considered in this experiment. As a first metric, we report the cross-entropy score on the latent space (CE, as in previous experiments). Then, we propose a comparison of the evidence lower bound (ELBO) on observed samples. Although in this experiment we do not have held-out data because we may only observe one cell at each leaf of the tree, we expect these numbers to be comparable because the same neural architecture, noise models, and hyperparameters were used for fitting both models. Our results, reported in Table 3, suggest that TreeVAE better fits the data and proposes an approximate posterior that is more reflective of the tree structure.

**Table 3:**
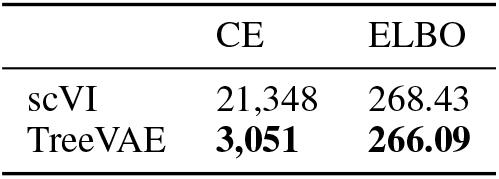
Results on the cancer metastasis data. ELBO denotes the evidence lowerbound on observed data.

|  | CE | ELBO |
| --- | --- | --- |
| scVI | 21,348 | 268.43 |
| TreeVAE | **3,051** | **266.09** |

Finally, we investigate the behavior of TreeVAE’s predictions of ancestral gene expression. As expected, TreeVAE’s imputations are more certain closer to the observations at the leaves and becomes more uncertain for nodes closer to the root (Figure 3a). We next predict the ancestral gene expression of *CEACAM5*, an important cell adhesion molecule that is associated with metastatic invasion [20]. We observe a predicted pattern that broadly agrees with tree structure and observations at the leaves (Figure 3b) and offers rich hypotheses on the subclonal dynamics of *CEACAM5* expression. Critically, because our Bayesian model quantifies uncertainty, we can directly evaluate the stability of a given hypothesis, unlike naive averaging. Overall, these results underscore the promise of the TreeVAE model for gene expression prediction.

**Figure 3:**
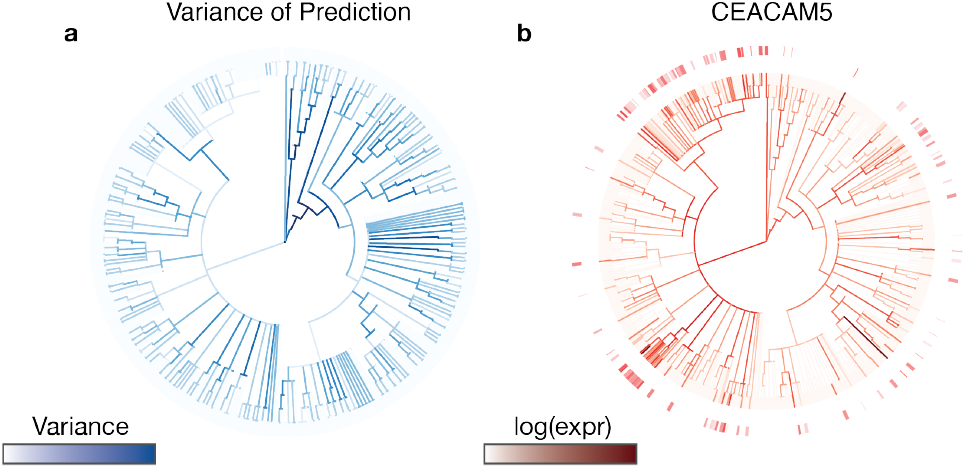
Behavior of TreeVAE internal node predictions. (a) Variance of posterior predictive density on latent space for each internal node. Uncertainty is negatively correlated with depth (Pearson’s *r* = −0.60). (b) Predicted expression of *CEACAM5* for each internal node. Color gradient at the leaves indicates observed gene expression.

## Discussion

By explicitly taking into account phylogenetic information for generative modeling of single-cell transcriptomic states, we introduce a framework for building more realistic latent representations of cells and inferring unobserved, or “ancestral”, intermediates. We demonstrate that TreeVAE provides robust predictions of unobserved cellular states and outperforms vector arithmetic in the latent space provided by state-of-the art algorithms like scVI [13]. Additionally, TreeVAE provides meaningful measures of uncertainty around its predictions, a critical feature for a researcher weighing several scientific hypotheses.

To the best of our knowledge, TreeVAE is the first generative model of single-cell measurements that takes into account a-priori dependencies between samples (cells). This is an under-explored area, as there are often technical challenges to fitting such models. The message-passing algorithm described in this manuscript, and the implementation available in PyTorch was sufficient for generating the results in this manuscript, although we are currently exploring several areas of future research around TreeVAE and similar models. First, we describe in this study a training procedure that uses full-batch updates. However, we anticipate that a more sophisticated, minibatch-based training scheme is necessary for improving scaling the inference procedure to larger trees (e.g., those with >50,000 cells). Second, the inference algorithm for TreeVAE could be improved by exploiting just-in-time compilation, for example with JAX [21], to speed-up the message passing algorithm.

We anticipate that the TreeVAE model described here will set the stage for several analytical avenues of interest. First, we expect that integrating accurate representations of unobserved cellular intermediates with accurate uncertainty quantification will enable a more sensitive differential expression analyses on the phylogeny. Second, the representations of ancestral states will also enable the identification of putative causal genes leading to particular phylodynamic patterns like subclonal expansions or differential metastatic behavior (as in [3]). Finally, these robust representations throughout a phylogenetic model will aid in the development of quantitative models of cell-state transitions throughout dynamic processes.

There is currently an unmet need to develop datasets that would aid in the construction and validation of models like TreeVAE. Ideally, these datasets would track at least a subset of marker genes dynamically over the course of development and could be used to evaluate predictive accuracy on real scRNA-seq data. Utilizing time-series datasets from model organisms for which cell lineages are stereotypical, like that of *C. elegans*, could be of great use towards this aim (e.g., [22]). Beyond this, we envision that maturing light-sheet microscopy technologies [23, 24] could be paired with unbiased readout of cellular transcriptomic states to create comprehensive molecular atlases of dynamic processes.

Since the first report of CRISPR/Cas9-based single-cell lineage tracing [25], the promise of these technologies has been to build complete, probabilistic fate maps of complex, dynamic biological processes. Yet, this promise has yet to be fully realized. Towards this, we foresee the TreeVAE model setting a foundation for this aim and beyond. With efforts around improving the engineering of the software, formulating sound statistical questions from these inferences, and developing sufficient ground truth datasets for evaluation we believe that more sophisticated TreeVAE models will be a key tool in the single-cell lineage tracing toolbox. Taken together, we proffer that continued development of such a tool would substantially work towards the ultimate promise of these lineage tracing technologies.

## Code Availability

The code to reproduce the experiments of this manuscript is available on GitHub https://github.com/khalilouardini/treeVAE-reproducibility. TreeVAE was implemented using an earlier version of the scvi-tools codebase [26].

## Acknowledgements

We first acknowledge members of the Yosef lab for helpful discussion. Additionally, we acknowledge Matthew Johnson and Sharad Vikram for their useful and interesting pointers on SVAEs and related topics.

## Competing Interests Statement

NY is an advisor and/or has equity in Celsius Therapeutics, and Rheos Medicine.

## Author Contributions

RL, MGJ, and NY conceived of the project and experiments, with input from SP and MIJ. KO and RL developed the message passing algorithm and training procedure, with input from MGJ. KO, RL and MGJ. wrote code for the TreeVAE model. RZ generated ground truth phylogenies for simulated experiments. KO performed all experiments with mentorship from RL, MGJ, and NY. MGJ performed analysis on the experimental metastasis data. NY supervised the completion of the work. All authors contributed towards writing the manuscript.

## A Message Passing Algorithms

In the main text of this manuscript, we presented a simple marginalization procedure provided for a triplet of nodes, based on completing the square. The procedure may be used recursively as part of a well-established message passing algorithm for computing marginal likelihood of leaf observations in the context of binary trees, and in particular explained in the TMC-VAE manuscript [10]. In this appendix, we present a generic message passing algorithm for multifurcating trees (Section A.1). In particular, we propose a generalization of the normalizing constant formula for multifurcating trees, a key contribution for dealing with phylogenetic information inferred from Cassiopeia and other algorithms that do not necessarily produce binary trees. We then demonstrate how to use this algorithm for computations of the marginal likelihood of the leaves *p*(*z*_1:*L*_) (Section A.2) and for posterior predictive densities *p*(*z_i_* | *z*_1:*L*_) (Section A.3).

### A.1 The base message passing algorithm

Let 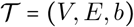 be a phylogeny with vertex set *V*, edge set *E* and edge length function *b*. We note the weight *b*(*e*) of an edge *e* = (*u, v*) as *b_u,v_*. Again, the vertex set *V* is partitioned into *L* leaf vertices *V_L_* = {1, …, *L*} (a set of cells) and *I* internal vertices *V_I_* = {*L* + 1, …, *L* + *I*} (ancestral cell states) such that *V* = *V_L_* ∪ *V_I_*. For a node *i* let *I* = *i*_1_, …, *i_n_* denote the indices of its n children.

Message passing is defined recursively, starting from a source node *s* (always an internal node), and requesting messages from each of its neighbors. We initialize the message of each the leaf *l* with the following content:

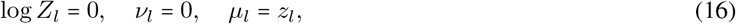

where *z_l_* is the evidence at leaf *l*.

Then, we use the following update rules to propagate messages from a set of child nodes (*i*_1_, …, *i_n_*) to a parent node *i*:

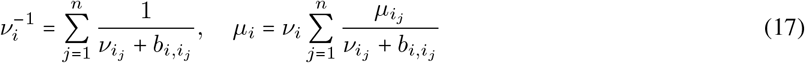

and for the normalizing constant:

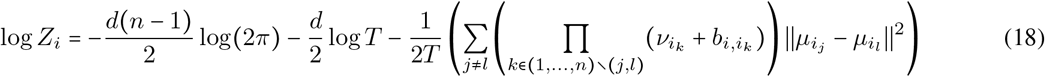

with 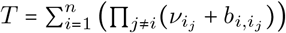.

### A.2 Computing marginals

In order to calculate the marginal likelihood of leaf observations *p*(*z*_1:*L*_), we run the message passing algorithm using the root node as the source, and using the leaf observations (*z*_1_, …, *z_L_*) as local evidence. This is described in [10], although our last steps differ from it, as they did not integrate out the prior in their calculations. For this, we compute the integral of the last message with the prior:

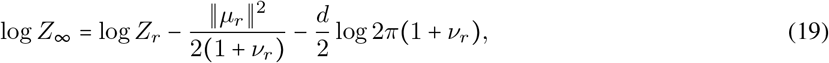

where *r* indicates the root. We then calculate the desired marginal distribution as:

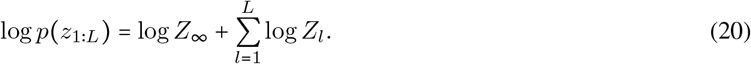

### A.3 Computing posterior predictive densities

In order to calculate the conditional distribution *p*(*z_i_* | *z*_1:*L*_) for an arbitrary internal node *i*, we run the message passing algorithm using the query node *i* as the source and using the leaf observations (*z*_1_, …, *z_L_*) as local evidence. In contrast to [10], we take into account the prior information during the message passing protocol. To do this, we add a dummy node attached to the root with null evidence and unit diagonal variance.

### A.4 Implementation and unit tests

We implemented this message passing information in vanilla PyTorch. In order to check our calculations, we have developed a suite of unit tests for random trees, with ground truth based on the Gaussian conditioning formula.

## B Gaussian Process Factor Analysis Model

In this appendix, we describe the Gaussian process factor analysis model used for simulating data, and define a tractable ground truth for both posterior predictive densities.

### B.1 Full specification of the generative model

Let *τ* = (*E, V, b*) be a phylogeny with *N* nodes. We index the vertex set 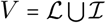 by leaves 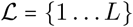 and internal nodes 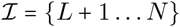. As for the TreeVAE model, latent variables *z_v_* for each vertex *v* form a multivariate Gaussian vector:

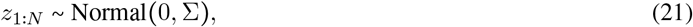

where Σ has for shape *Nd* × *Nd* (the latent space is *d*-dimensional). For every leaf *n*, observation *x_n_* is generated as:

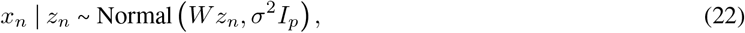

with *W* ∊ ℝ^*p*×*d*^ and *σ* > 0 (the feature space is *p*-dimensional).

### B.2 Posterior distribution

The posterior distribution *p*(*z*_1:*L*_ | *x*_1:*L*_) is tractable via Gaussian conditioning formula. Indeed, it is easy to see that (*z*_1:*L*_; *x*_1:*L*_) is a Gaussian vector:

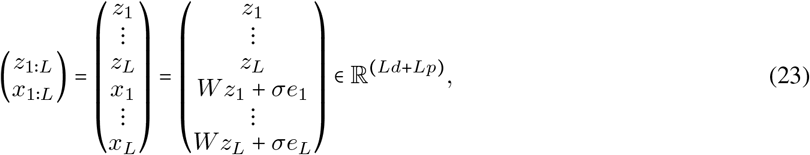

where *e*_1:*L*_ is sampled from a Gaussian isotropic distribution. Consequently, we can characterize the distribution of this vector by its mean and covariance. The random vector is centered, as *z*_1:*L*_ and *e*_1:*L*_ are both centered. Let us denote the covariance matrix of (*z*_1:*L*_, *x*_1:*L*_) by Λ ∊ ℝ^(*Ld*+*Lp*)×(*Ld*+*Lp*)^. We decompose Λ with a block structure:

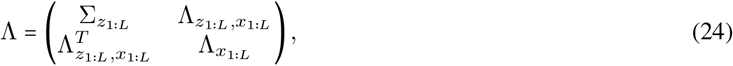

and where each term can be calculated as follows:

#### Marginalized latent covariance

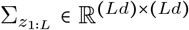 the marginalized covariance Σ of the leaves (computed by adequately slicing Σ).

#### Marginalized feature covariance

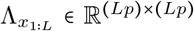 such that for all pairs of leaves 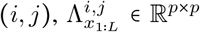 is the block encoding correlations between *x_i_* and *x_j_*:

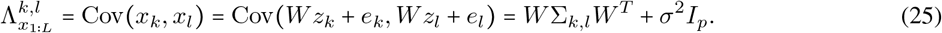

#### Correlation term

The matrix 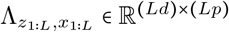 encodes correlations between *z*_1:*L*_ and *x*_1:*L*_ such that:

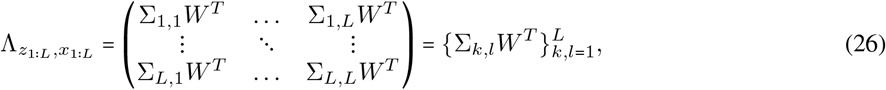

where the previous result comes from the simple fact that for a pair of leaves (*k, l*), we have:

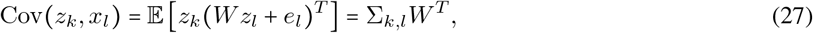

where Σ*_k,l_* is the marginalized Σ corresponding to the correlations between variables *z_k_* and *z_l_*.

Finally, we may use the Gaussian conditioning formulas to derive the posterior distribution *p*(*z*_1:*L*_ | *x*_1:*L*_):

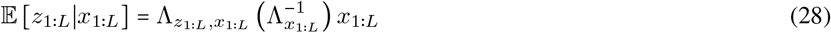

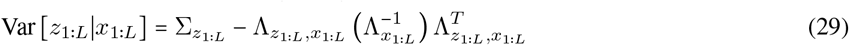

### B.3 Posterior predictive densities at internal nodes

In all our imputation experiments, we are interested in the posterior predictive density of the internal nodes. For both quantities, we integrate over a suitable set of latent variables. On latent space, we utilize the following decomposition:

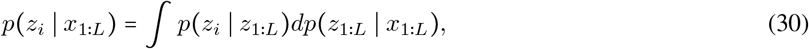

where *p*(*z_i_* | *z*_1_ … *z_L_*) may be computed exactly using the message passing algorithm (as in Section A.3) for any internal node *i*. On feature space, we similarly proceed and integrate out latent variables:

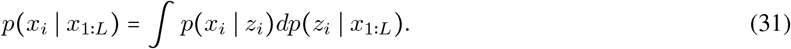

Therefore, we can efficiently compute the posterior predictive of any internal node through MCMC sampling in two steps. First, we sample from the posterior predictive distribution on latent space at node *i*: *p*(*z_i_* | *x*_1:*L*_). Then, we generate *x_i_* | *z_i_* with the generative model specified in Section B.1.

## Notes

https://github.com/khalilouardini/treeVAE-reproducibility

### Competing Interest Statement

NY is an advisor and/or has equity in Cellarity, Celsius Therapeutics, and Rheos Medicine.

